# Emotional state dynamics impacts temporal memory

**DOI:** 10.1101/2023.07.25.550412

**Authors:** Jingyi Wang, Regina C. Lapate

## Abstract

Emotional fluctuations are ubiquitous in everyday life, but precisely how they sculpt the temporal organization of memories remains unclear. Here, we designed a novel task—the Emotion Boundary Task—wherein participants viewed sequences of negative and neutral images surrounded by a color border. We manipulated perceptual context (border color), emotional valence, as well as the direction of emotional-valence shifts (i.e., shifts from neutral-to-negative and negative-to-neutral events) to create encoding events comprised of image sequences with a shared perceptual and/or emotional context. We measured memory for temporal order and subjectively remembered temporal distances for images processed *within* and *across* events. Negative images processed within events were remembered as closer in time compared to neutral ones. In contrast, temporal distance was remembered as longer for images spanning neutral-to-negative shifts— suggesting temporal dilation in memory with the onset of a negative event following a previously-neutral state. The extent of this negative-picture induced temporal dilation in memory correlated with dispositional negativity across individuals. Lastly, temporal order memory was enhanced for recently presented negative (compared to neutral) images. These findings suggest that emotional-state dynamics matters when considering emotion-temporal memory interactions: While persistent negative events may compress subjectively remembered time, dynamic shifts from neutral to negative events produce temporal dilation in memory, which may be relevant for adaptive emotional functioning.

## Introduction

The seemingly continuous unfolding of our lives can be rippled by internal and external changes. Prior studies indicate that acute contextual shifts—such as going from one room to another—form so-called event boundaries (Zacks, 2020), which sculpt the organization of episodic memories, including memory for their temporal features (temporal memory) (Clewett et al., 2019; Shin & DuBrow, 2021). For example, subjective estimates of the time elapsed between a pair of items (henceforth referred to as ‘temporal distance’) are longer when items span an event boundary (even when objective time is held constant) (Ezzyat & Davachi, 2014; McClay et al., 2023; van de Ven et al., 2022; Y. C. Wang & Egner, 2022; Wen & Egner, 2022). Moreover, memory for the order in which items were originally encoded (henceforth referred to ‘temporal order memory’) is often impaired for items spanning an event boundary (Clewett & Davachi, 2021; DuBrow & Davachi, 2014, 2016; Gurguryan et al., 2021; Heusser et al., 2018; Hsieh et al., 2014; McClay et al., 2023; Pu et al., 2022; Sols et al., 2017; van de Ven et al., 2022; Y. C. Wang & Egner, 2022; Wen & Egner, 2022). These contextual shifts that form event boundaries are often accompanied by internal emotional-state changes, such as the shift to a joyful mood upon opening the lab door where a surprise birthday party awaits. Nonetheless, despite the ubiquity of internal, emotional-state changes as we navigate shifting external contexts in our everyday lives, whether and how dynamic emotional states shape temporal memory—potentially impacting event boundaries produced by external contextual changes—has only recently been studied (Clewett & Davachi, 2021; Dev et al., 2022; McClay et al., 2023; Palombo et al., 2021).

It is well established that high-arousal negative items are better remembered than neutral ones (Bowen et al., 2018; Cahill & McGaugh, 1995; Kensinger & Corkin, 2003; LaBar & Cabeza, 2006) but prior work suggests that memory for contextual, episodic details surrounding emotionally evocative stimuli is not always facilitated by emotion, and may be deprioritized instead (Bisby & Burgess, 2017; Cohen & Kahana, 2022; Kensinger, 2009; Mather, 2007; Mather & Sutherland, 2011; Palombo & Cocquyt, 2020; Rimmele et al., 2011; Talmi et al., 2019). Prior studies have revealed privileged recall of emotional content, at times at the expense of temporally organized retrieval (Barnacle et al., 2016; Long et al., 2015; Siddiqui & Unsworth, 2011)—a phenomenon that is captured by recent computational models detailing mechanisms of disruption of temporal context encoding by emotional items (Cohen & Kahana, 2022; Talmi et al., 2019). Consistently, new anatomical and functional evidence suggests that high-arousal negative events may impair memory for temporal context via amygdala-hippocampal interactions and local competition (J. Wang et al., 2022). Accordingly, picture-induced negative emotion can produce sustained functional coupling in amygdala-hippocampal circuitry and hippocampal neural activity patterns for tens of minutes, which carries over into epochs of novel neutral-stimulus encoding (Tambini et al., 2017)—an effect specifically observed following negative-to-neutral transitions (compared to neutral-to-negative transitions). Further underscoring the pervasiveness of negative affect, prior work indicates that negative emotional processing can persist beyond its temporal context to bias appraisals of novel neutral stimuli, suggesting a potential blurring of event boundaries by negative emotional experiences (Lapate et al., 2016, 2017). Collectively, this work suggests that negative emotional processing may impact the processing of temporal context during and/or following the onset of negative events.

Results from existing behavioral studies have been mixed, and indicate that salient, negative emotions may differentially impact specific aspects of temporal memory. For example, studies examining how negative emotional processing modulates temporal *order* memory have reported both impairments (Huntjens et al., 2015; Maddock & Frein, 2009; Zlomuzica et al., 2016) and enhancements (D’Argembeau & Van der Linden, 2005; Dev et al., 2022; Rimmele et al., 2012; Schmidt et al., 2011). On occasion, emotional valence shifts may improve temporal order accuracy while distorting temporal distance judgments. For instance, a recent study using music to induce emotion during neutral object encoding showed that shifts towards positively valenced states strengthened temporal order memory while shortening the remembered temporal distance between neutral object pairs (McClay et al., 2023). Findings from studies examining how emotional events impact retrospective temporal duration judgments align with this result, and suggest temporal dilation for negative compared to neutral events (M. Anderson et al., 2007; Bisson et al., 2008; Campbell & Bryant, 2007; Johnson & MacKay, 2019; Loftus et al., 1987; Stetson et al., 2007).

Nonetheless, prior work has only rarely explicitly modeled the impact of negative emotion on temporal memory for items encoded during prolonged negative events compared to following transitions *to* and *from* negative emotional events. This is important, as prior neural and behavioral work shows that negative emotional provocations often linger onto neutral-event epochs (but not vice-versa) (e.g., (Tambini et al., 2017)). Moreover, recent work suggests that distinct emotional-state transitions—such as from positive-to-negative, and from negative-to-positive states—can exert distinct impacts on temporal memory—i.e., subjective temporal dilation vs. compression^1^, respectively (McClay et al., 2023). However, available studies that have considered directional emotional-event shifts (produced by music) or shifts in mildly arousing stimuli (produced by neutral tones) have thus far examined temporal memory for (unrelated) neutral items presented concurrently with the emotion-inducing stimuli (Clewett & Davachi, 2021; McClay et al., 2023)—as opposed to temporal memory for the emotional events themselves. This is important, as features that are inherent or intrinsic (vs. extrinsic) to an emotional event may be differentially prioritized in memory (Anderson & Shimamura, 2005; Bisby & Burgess, 2013, 2017; Davachi & DuBrow, 2015; Kensinger, 2009; Kensinger et al., 2007; Kim et al., 2013; Lee et al., 2014; Mather, 2007; Ventura-Bort et al., 2016). Relatedly, emotional events may occur synchronously or asynchronously with other external, contextual shifts, which themselves influence temporal memory (Peterson et al., 2021). Together, these open questions require examining whether and how dynamic, emotional events shape temporal memory for those events—and whether changes in other contextual (e.g., perceptual) features interact with emotional changes to sculpt temporal memory.

Finally, there remains a suggestive but little understood association between emotional processing, temporal memory, and well-being. Altered temporal dynamics of emotional responding, including the temporal persistence of negative affect over time, are ahallmark of mood disorders (Davidson, 2000; Lapate & Heller, 2020; Puccetti et al., 2021;Schaefer et al., 2013). Atypical temporal memory in mood and anxiety disorders has alsobeen noted. For instance, individuals with depression often omit time-related descriptors(e.g., “now”, “later”) in their narratives, and tend to recall autobiographic memories in a non-linear fashion compared to healthy controls (Habermas et al., 2008). Similarly, patients diagnosed with post-traumatic stress disorder (PTSD) usually show temporally disorganized recall of traumatic events (Ehlers & Clark, 2000). When asked to sort negatively provocative images into their originally presented order, lower temporal order accuracy has been found in individuals with higher anxiety (Huntjens et al., 2015). Collectively, these findings raise the possibility that individual differences in memory for temporal contexts processed during or following negative emotional events may be related to vulnerability to psychopathology.

Therefore, the goal of the present study was to investigate how high-arousal, negative emotional events—and accompanying dynamic emotional-event transitions—shape temporal memory for those emotional events, including following external changes in context. To that end, we designed a novel task, the ***Emotion Boundary Task***, which employed sequences of negative and neutral images presented at fixed time intervals, surrounded by a color border. By manipulating the emotional valence^2^ of sequentially presented pictures, as well as the direction of emotional-valence shifts, we produced two emotional event types (Negative and Neutral) and two distinct kinds of “emotion boundaries” (from Neutral-to-Negative, and from Negative-to-Neutral). These emotional events and emotional-event transitions were orthogonally manipulated in relation to changes in the color of a border surrounding the pictures, a visuo-perceptual change known to produce event boundaries (Heusser et al., 2018; Wen & Egner, 2022). Thus, we produced encoding events comprised of image sequences that shared the same border color and/or emotional valence. Following the presentation of each encoding list of the Emotion Boundary Task, we measured subjectively remembered temporal distances^3^ as well as temporal order memory for pairs of images sampled from within and across boundary conditions (**Figure 1**). This allowed us to examine how emotional events and directional changes between those events modulate memory for temporal distance and order—as well as whether these emotional factors interact with perceptually-originated contextual shifts (produced by changes in border color). We hypothesized that negative to neutral event transitions might blur event boundaries, reducing temporal distance estimates (J. Wang et al., 2022; Zlomuzica et al., 2016), whereas neutral to negative event transitions may dilate temporal distance estimates and potentially increase temporal order memory accuracy (Dev et al., 2022; Johnson & MacKay, 2019); see also (McClay et al. 2023). Finally, we examined whether temporal memory distortions following negative emotional-event shifts would be associated with individual differences in affective style linked to psychopathology risk (Shackman et al., 2016).

**Figure 1.**
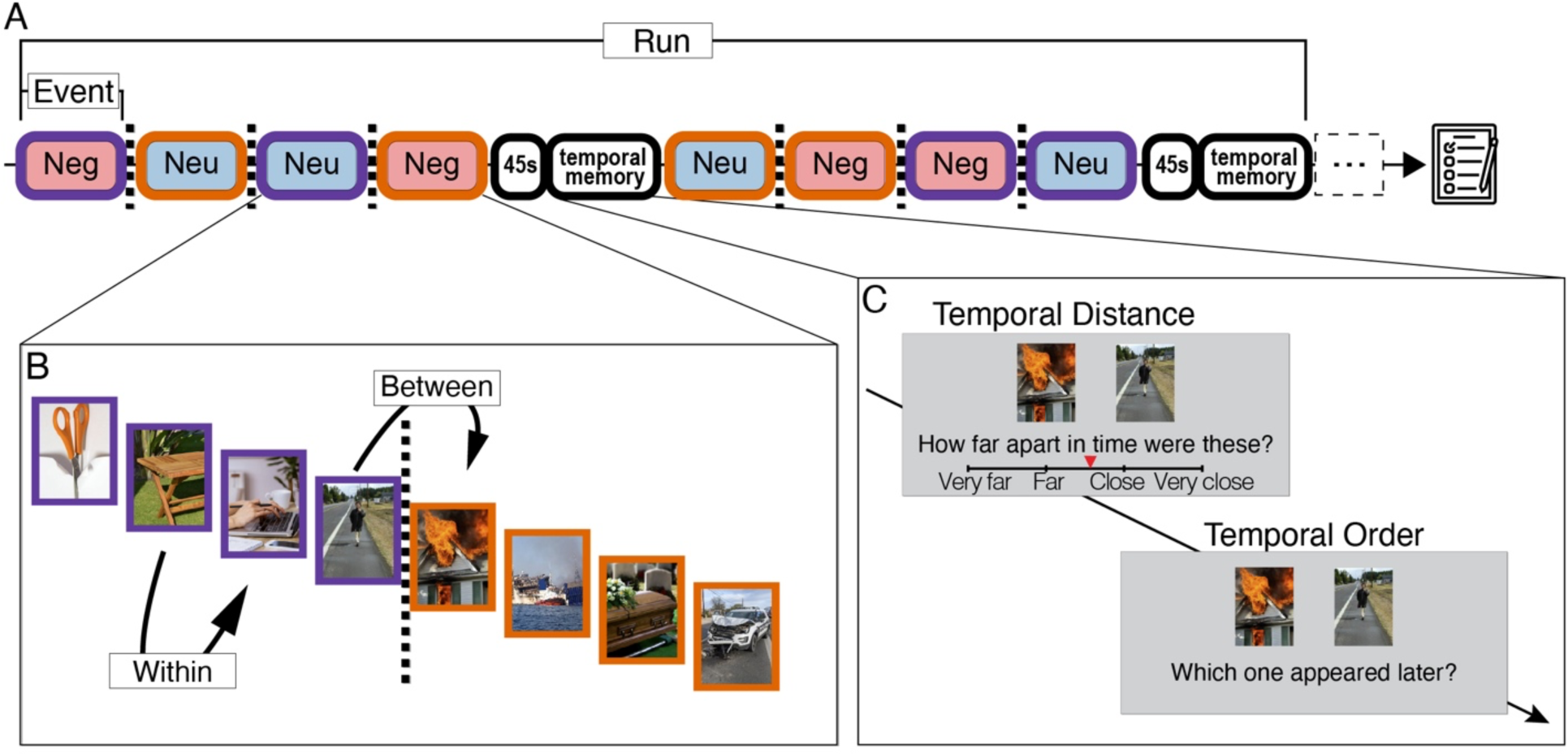
Schematic of the emotion-boundary and temporal memory tasks. **(A)** The Experiment Timeline. In the Emotion Boundary Task, participants studied sequences of emotional and neutral images, which were surrounded by a color border. Each event consisted of four sequential images displaying a particular emotional valence and/or border color. Event boundaries (dotted lines) were created by changing image valence, border color, or both. After encoding four events (16 images total), participants performed a 45-s distractor task, followed by temporal memory tasks (assessing temporal distance and order memory). At the end of the experiment, participants completed mood questionnaires. Dotted lines indicate event boundaries. Red indicates negative valence (‘Neg’). Blue indicates neutral valence (‘Neu’). **(B)** Two event sequences are shown. **(C)** Temporal distance and order memory tasks are shown. Pairs of images were sampled from within and between-event conditions for testing. Participants were instructed to indicate remembered (subjective) temporal distances between item pairs as well as the presentation order of each image pair.

## Materials and methods

### Participants

Eighty-two healthy young adults (23 males; mean age = 20.07 years, age range = 18-26 years) were recruited from the University of California, Santa Barbara. Two participants were excluded due to poor quality data (see *Participants Exclusion* under *Statistical Analysis*). The remaining sample (N = 80; 23 males; mean age = 20.1 years, age range = 18-26 years) was used for all analyses. Due to COVID-19 related University closures (2020-2021), data were collected online using Pavlovia (task data; https://pavlovia.org) and Qualtrics (questionnaire data; https://www.qualtrics.com/). All participants signed a consent form prior to undergoing the study. All study procedures were approved by the Institutional Review Board at the University of California, Santa Barbara. All task and analysis code can be found at https://osf.io/zr7hx/.

### Stimuli

A total of n = 192 pictures were selected from the International Affective Picture System (Lang et al., 2008). Using the IAPS’ standardized ratings for emotional valence and arousal (where 1 = very negative/low-arousal and 9 = very positive/high-arousal), we selected 96 negative images (M_valence_ = 2.23, *SD* = 0.33; M_arousal_= 5.89, *SD* = 0.71) and 96 neutral images (M_valence_ = 5.10, *SD* = 0.28; M_arousal_= 2.93, *SD* = 0.38). As expected, negative images were significantly more negative and arousing than neutral images (t_(184.45)_ = 65.57, *p* < 0.001, and t_(144.96)_ = −36.10, *p* < 0.001, respectively).

## Procedure

### Overview

To determine the impact of emotional state changes on temporal memory, we manipulated the emotional valence of sequential negative and neutral images shown in an event-boundary paradigm (Clewett & Davachi, 2021; Heusser et al., 2018), which we term the *Emotion Boundary Task*. In this task, we orthogonally manipulated border color change (Change, Non-change), the valence of the first image in the tested pair (Neutral, Negative), and the valence of the second image in the tested pair (Neutral, Negative) – thereby yielding two directions of emotional-event shifts (Negative-to-Neutral and Neutral-to-Negative). Therefore, these factors yielded unique event-boundary conditions as a function of emotional valence, border color, and emotional-shift direction (dotted lines in **Figure 1**). Image pairs from within and across these perceptual and emotion boundaries were sampled for temporal memory assessments, which were interleaved with the encoding task. At the end of the experiment, participants completed trait mood and anxiety questionnaires. As part of a larger study, participants completed additional temporal processing and memory tasks at the end of the experiment (data not reported here).

### Emotion Boundary Task

In each task block, participants were shown unique negative and neutral images surrounded by a color border (orange or purple). Following prior work, to promote continuous task engagement, participants were instructed to judge each image on whether they expected to see the current border color in it (‘yes’ or ‘no’) (DuBrow & Davachi, 2013, 2014, 2016; Ezzyat & Davachi, 2014; Gurguryan et al., 2021; Heusser et al., 2018; Pu et al., 2022; van de Ven et al., 2022; Wen & Egner, 2022). Participants were asked to respond as fast as possible upon seeing each image by pressing one of two keys (‘left arrow’ versus ‘right arrow’). Each image was shown for 4 s, followed by a 2.5 s intertrial interval (ITI). A fixation cross was displayed continuously at the center of the screen, and participants were instructed to maintain central fixation throughout the task.

Four sequential images (displaying the same emotional valence and border color) comprise what we term an ‘event’. The color border remained visible throughout each event. Following prior work, changes in image border color and emotional valence across events were expected to serve as event boundaries and/or emotion boundaries (DuBrow & Davachi, 2013, 2014, 2016; Ezzyat & Davachi, 2014; Heusser et al., 2018).

Because a primary goal of the current study was to specify whether the *direction* of emotional-state transitions modulated temporal memory, we generated an equal number of image sequence transitions from Negative-to-Neutral and Neutral-to-Negative. Image pairs were sampled from within and across events for subsequent temporal memory testing, which included equal sampling of negative and neutral images shown earlier versus later within a particular encoding sequence—hereafter, *First and Second Image(s)—*thereby yielding a 2 x 2 x 2 design: First Image Emotional Valence (2: Neutral, Negative); Second Image Emotional Valence (2: Neutral, Negative); and Change in Image Border Color (2: Change, no Change).

After every set of four events (referred to here as ‘encoding list’), a 45-s distractor task began, which was followed by temporal memory assessments (**Figure 1**). Each task run comprised two encoding lists, which including one instance of every boundary transition type (**Supplementary Figure 1**), after which participants were offered a short break. A total of six unique event-boundary runs were shown to each participant. Image order as well as the order of event-boundary runs were randomized across participants. Inaddition, we counterbalanced the occurrence of negative and neutral events at both the first and last position across runs.

### Distractor Task

To attenuate recency effects, participants performed a distractor task for 45s after each set of four events (i.e., 16 images), before undergoing temporal memory assessments (**Figure 1**). In this distractor task, a white circle randomly moved either to the right or to the left. Participants’ task was to hold the right (vs. left) arrow key to follow the circle trajectory as it moved to the right (vs. left).

### Temporal Memory Tasks

After the 45-s distractor task, participants were presented with pairs of images they had previously seen, and asked to provide a subjective temporal distance judgment (“*How far apart in time were these two images?*’) and judge the order in which they were originally presented in the sequence (“*Which image appeared later in the sequence?*’) (**Figure 1C**). We instructed participants to indicate remembered temporal distances by stating *“We will ask you to try to guess the time interval between two images*”, alongside an example of the temporal distance task slide during the initial task overview. Participants rated the remembered temporal distance between each pair of images by placing a marker on a continuous slider, which was anchored by four tick marks (“very far”, “far”, “close” and “very close”) from left to right. The slider range was [1-4], in which “very close” was coded as 1, “close” was coded as 2, “far” was coded as 3, and “very far” was coded as 4. Following prior work (Wen and Egner, 2022; Wang and Egner, 2022; McClay et al., 2023), the marker was set to the middle of the slider at the beginning of each temporal distance trial. Given that the objective time intervals between tested image pairs were fixed (Clewett et al., 2020; Clewett & Davachi, 2021; Heusser et al., 2018; Pu et al., 2022; Sols et al., 2017; van de Ven et al., 2022; Wang & Egner, 2022; Wen & Egner, 2022), observed differences in the temporal distance metric reflect *subjective* time—as remembered by participants. Next, participants indicated the temporal order in which the same pair of images was shown by pressing the left or right arrow key to select the image they thought had been presented more recently.

Within-event and between-event image pairs for subsequent temporal memory testing were selected from each encoding list (Figure 1 and Supplementary Figure 1). Within-event pairs comprised the second and third images presented within an event, whereas between-event pairs were comprised of fourth image from one event and the first image from the subsequent event. Seven image pairs (4 within-event and 3 between-event) constituted all adjacent pairs of images to be tested per encoding list (see *Supplementary Figure 1* for an illustration of item pairs tested for a given encoding sequence). The screen location of each image in a pair (right versus left side of the screen) was randomized. Temporal memory test trials were self-paced, followed by a 0.5 s ITI.

### Mood questionnaires

At the end of the experiment, participants completed the following self-reported mood questionnaires: the *Beck’s Depression Inventory* (Beck et al., 1961), the *State-Trait Anxiety Inventory* (Spielberger et al., 1983), and the trait *Negative Affect* (NA) subscale from the short-form of the *Adult Temperament Questionnaire* (Evans & Rothbart, 2007). Using the data obtained from these questionnaires, we computed a *dispositional negativity* factor score—a facet of trait-like affective style reliably implicated in psychopathology vulnerability (Shackman et al., 2016)—for each participant. To do so, we performed a principal component analysis (PCA) on the questionnaire data [principal function; Psych R package (Revelle, 2022)]. Together, the three questionnaires explained 65.4% of the variance in the dispositional negativity score (λ_BDI_ = 0.86, λ_STAI_ = 0.95, λ_ATQ_ = 0.56). The resulting dispositional negativity scores were correlated with emotion-driven changes in temporal memory. Because the loading of ATQ in the dispositional negativity factor was unexpectedly low, we additionally examined the association between trait emotionality and temporal memory using dispositional negativity scores obtained using the BDI and STAI-Trait only (88.9% of the variance explained; λ_BDI_ = 0.94, λ_STAI_ = 0.94)— which replicated the results from our a-priori (3-questionnaire) factor analytical approach.

### Statistical analysis

#### Participant exclusion

Throughout the task, participants were asked to indicate whether they expected to see the border color in each image shown to them (‘yes’ or ‘no’). We used this color-image encoding task to encourage participants to pay attentionto each image and its border, and to monitor participant engagement throughout the task(Heusser et al., 2018). Two participants (2/82; 2.4%) were excluded due to low responserates (i.e., response omission in > 25% of all trials).

#### Mixed-effects modeling

To examine the impact of emotional events and emotional-event shifts on temporal distance and order memory, we modeled the temporal distance data using mixed-effects linear regression models and the temporal order memory data using mixed effects logistic regression in R (Bates et al., 2015). Since all factors manipulated were nested within participants, mixed-effects models were fit with random factor slopes and random intercepts per participant. When appropriate, pairwise comparisons were run in R (Lenth, 2021). Bayes factors for mixed-effect models were calculated following Dienes, 2014 and 2019 (https://github.com/HTattanBirch/bayes-factor-calculator).

#### Correlation between temporal distance scores and dispositional negativity

As detailed in the Results, we observed a robust impact of Neutral-to-Negative event transitions on temporal distance ratings, which were consistent with a temporal dilation effect. To examine whether individual differences in the magnitude of this emotion-driven change in temporal memory were associated with individual differences in affective style, we averaged temporal distance scores reflecting the impact of emotional state shifts we observed (i.e., temporal distance dilation following *Neutral-to-Negative* events) per participant. We focused on emotional-event shifts obtained within color event boundaries (as opposed to across color events) to obtain scores reflecting emotion-driven changes in temporal distance without the additional known impact of border color changes on temporal memory (DuBrow & Davachi, 2014, 2016; Ezzyat & Davachi, 2014; Wen & Egner, 2022). To examine effects obtained during the transition from Neutral-to-Negative events while controlling for inter-individual variation attributable to other non-specific effects, we performed a residualization procedure using multiple regression, whereby we regressed [Negative-to-Neutral] on [Neutral-to-Negative] scores and saved the residuals. In other words, this residualized score reflects inter-individual variation in [Neutral-to-Negative] event transitions while controlling for inter-individual variation in other non-specific factors (e.g., rating biases, overall memory, etc.). This residualization procedure maximizes the reliability of correlations between changes in temporal distance ratings and other metrics as well as avoids the complexity of interpreting results from double difference scores (Lapate et al., 2017; Tucker et al., 1966). To ascertain emotion condition specificity, we performed the analogous residualization procedure for [Negative-to-Neutral] scores (controlling for [Neutral-to-Negative] scores). These residualized scores were then separately correlated with dispositional negativity using Pearson’s correlation. Of note, this approach is equivalent to that of conducting a multiple regression that includes temporal distance scores obtained from the [Negative-to-Neutral] condition in a model that predicts dispositional negativity from [Neutral-to-Negative] temporal distance scores. Next, using the Cocor package (Diedenhofen & Musch, 2015), we tested the difference in correlation coefficients between the Pearson’s correlation of dispositional negativity and temporal distance memory obtained for distinct event boundary conditions. ***Correlation between temporal distance and order memory.*** To examine whether there was a relationship between emotion-driven changes in temporal distance and temporal order memory, we averaged temporal distance ratings and temporal order accuracy per participant (within border color events). Then, we calculated difference (delta) scores between every condition pair for each type of temporal memory (**Table 1**). We correlated the resulting temporal distance and temporal order accuracy delta scores using Pearson’s correlation.

**Table 1.** Correlation between temporal distance and order metrics.

| **Condition pairs** | **Pearson's r** | **p** |
| --- | --- | --- |
| within Neutral – within Negative | 0.07 | 0.55 |
| Neutral-to-Negative – Negative-to-Neutral | 0.01 | 0.90 |
| Neutral-to-Negative – within Neutral | 0.14 | 0.23 |
| Neutral-to-Negative – within Negative | 0.13 | 0.24 |
| Negative-to-Neutral – within Neutral | 0.10 | 0.36 |
| Negative-to-Neutral – within Negative | -0.19 | 0.09 |

#### Control Analyses

*Semantic similarity scores*. Prior work has suggested that semantic similarity can differ between negative vs. neutral emotional valences (Riberto et al., 2019, 2022)—and that these factors therefore could potentially confound analysis of emotion-related changes in temporal distance. To estimate the semantic similarity between all pair of images used in our study, we employed two deep convolutional artificial neural networks (ANNs): CORnet-S (Kubilius et al., 2018), and vgg-16 (Simonyan & Zisserman, 2014). Previous studies indicate that artificial neural activation patterns in the “IT” layer of the CORnet-S network and the “fc2” layer of the vgg-16 networks show good agreement with neural activations observed in the inferior temporal cortex, a brain area known to represent objects and to track semantic judgments (Charest et al., 2014; Clarke & Tyler, 2014; Dunsmoor et al., 2014; Haxby et al., 2014; Iordan et al., 2015; Kriegeskorte et al., 2008; Kubilius et al., 2018, 2019; Martin et al., 1996; Riberto et al., 2019; Schrimpf, et al., 2020; Tang et al., 2018). Therefore, we obtained the ANN-drived IT layer activation for each image using the “candidate_models” package (https://github.com/brain-score/candidate_models), which yielded an activation matrix for each image. Following prior work, a principal component analysis (PCA) was used to reduce the dimensionality of the data (Schrimpf, et al., 2020). Next, we calculated pairwise image semantic similarity scores by obtaining the Pearson’s correlation between the activation vectors of all pairs of images used in the study. To increase the reliability of this metric, we average the results obtained across two previously-validated neural networks (“CORnet-S”, “vgg16”), which showed good inter-network agreement (r_(18334)_ = 0.602, p < 0.001) (henceforth, this average score is referred to as “semantic similarity”). Finally, we entered semantic similarity scores as a covariate in the above-described mixed effects models examining the influence of perceptual and emotional valence shifts on remembered temporal distances.

## Results

### Dynamic emotional state changes modulate memory for temporal distance

Event boundaries produced by changes in visuo-perceptual cues (such as border color) often lengthen the remembered temporal distance for items processed across boundaries (DuBrow & Davachi, 2014, 2016; Ezzyat & Davachi, 2014; Wen & Egner, 2022). Consistently, we found that a change in border color produced an event boundary in our paradigm, significantly increasing remembered temporal distance for items spanning across (compared to within) color change events (main effect of border color, B = 0.059 (SE = 0.011), *F* = 27.119, *p* < 0.001, η^2^_p_ = 0.17, BF= 2.933×10^5^, **Figure 2A**).

**Figure 2.**
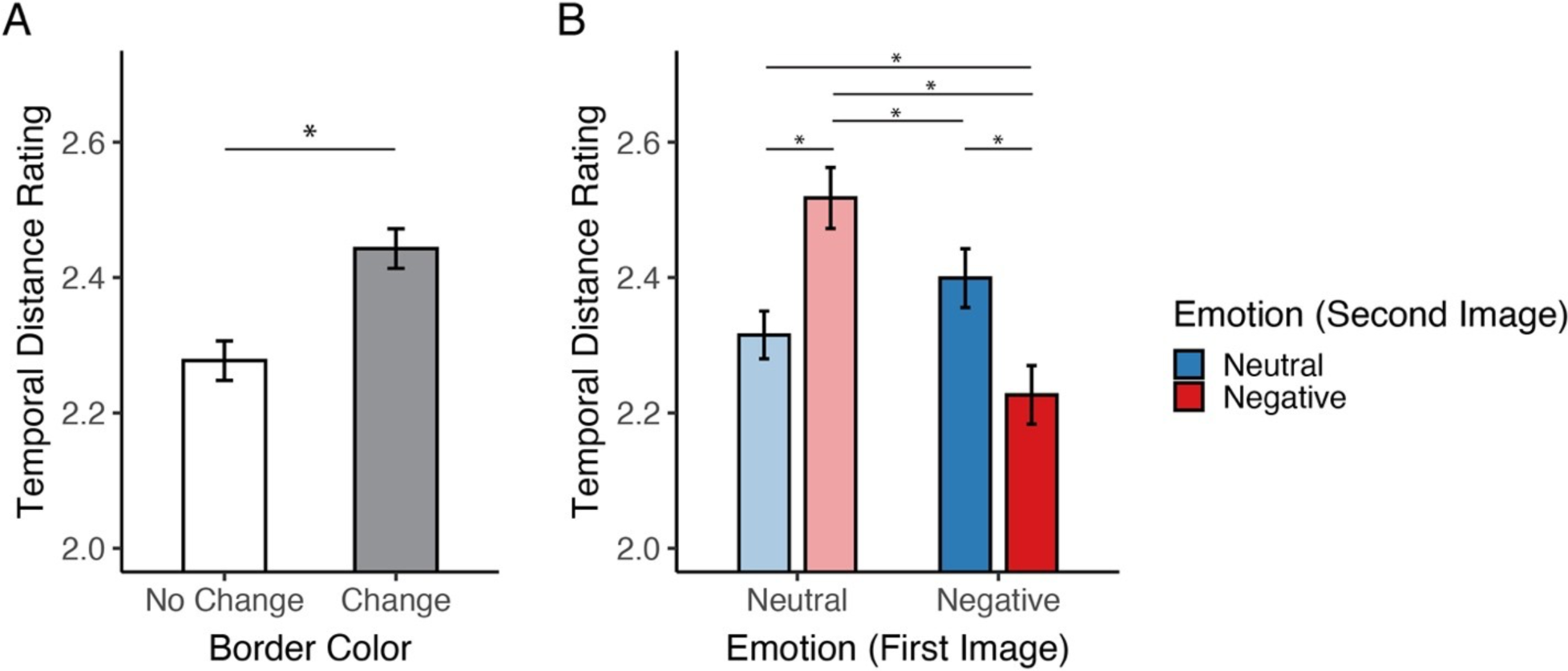
Impact of emotion and emotional-event transitions on temporal distance memory. **(A)** Temporal distances were remembered as longer for pairs of items sampled from across (vs. within) border color change events. **(B)** Emotional valence from the images encoded first and second from a tested image pair interacted in sculpting temporal distance memory, underscoring the directional impact of emotional-event shifts on temporal memory. Specifically, Neutral-to-Negative emotional-event transitions produced subjective temporal dilation in memory, whereas subjectively remembered temporal distances were compressed following sequential negative (vs. neutral) images. Error bars: within-subjects 95% confidence interval for each condition, computed using Morey’s (2008) method. \**p* < 0.05.

Next, we tested whether emotional valence—as well as the direction of emotional-valence shifts—influenced temporal distance memory. We found that the emotional valence of tested images shown earlier in the sequence modulated subjectively remembered temporal distances—such that whenever a negative image was shown first, the remembered temporal distance between it and the second image of the tested pair decreased—i.e., it was remembered as shorter—compared to when a neutral image was shown first (mixed effects model B = −0.070 (SE = 0.018), *F* = 14.630, *p* < 0.001, η^2^_p_ = 0.14, BF= 678.91). Critically, there was a significant interaction between the factors of first-image valence and second-image valence (B = −0.149 (SE = 0.023), *F* = 40.568, *p* < 0.001, η^2^ = 0.31, BF= 1.943×10^11^, see **Figure 2B**). This result indicates that the *direction* of emotional-valence shifts robustly modulated temporal distance memory. Specifically, shifting from Neutral-to-Negative events *increased* temporal distance ratings compared to Negative-to-Neutral shifts (*p* = 0.001, *d_z_* = 0.373). Consistently, the remembered temporal distance between images spanning Neutral-to-Negative transitions was larger than between image pairs shown within neutral (*p* < 0.001, *d_z_* = 0.775) or within negative events (*p* < 0.001, *d_z_* = 0.923). Of note, remembered temporal distances for negative images shown within negative events decreased—i.e., were remembered as shorter— compared to neutral images from within neutral events (*p* = 0.018, *d_z_* = 0.328) (**Figure 2B**). The three-way interaction of border color, first and second valence on temporal distance was not significant (B = 0.010 (SE = 0.02), F = 0.226, *p* = 0.635, η^2^_p_ = 0.001, BF= 0.06). (For full model results, including Bayes Factors, and means and SEMs per condition, see **Supplementary Figure 2** and **Supplementary Tables 1 and 2**). Collectively, these results suggest relative temporal compression in memory when processing sequential negative events, but temporal dilation when transitioning from neutral to negative events.

#### Control Analysis: Semantic Similarity

Because negative stimuli may have stronger semantic links than neutral stimuli, which could themselves drive participants’ temporal distance estimates (Dunsmoor et al., 2014; Riberto et al., 2019, 2022; Talmi, 2013), we computed semantic similarity scores for all image pairs used in this study, and re-ran our temporal distance mixed-effect model after entering them as a covariate (see *Methods*).

First, we found that semantic similarity was higher for image pairs with the same valence compared to those with different valences (t_(17623)_ = 23.430, *p* < 0.001); moreover, similarity was higher for negative image pairs compared to neutral pairs (t_(9098.9)_ = 2.481, *p* = 0.013), consistent with recent work (Riberto et al., 2019, 2022). In addition, similarity scores correlated negatively with temporal distance judgments (B = −0.526 (SE = 0.114), F = 21.359, *p* < 0.001, η^2^_p_ = 0.003, *r*_(6087)_ = −0.060, *p <* 0.001), suggesting that image similarity itself can influence temporal memory. Importantly, the impact of emotional valence transitions on temporal distance remained significant after accounting for pairwise image similarity by entering it as a covariate in a mixed-effects model. After controlling for semantic similarity, changes in border color continued to increase remembered temporal distances (B = 0.056 (SE = 0.014), F = 24.283, p < 0.001, η^2^_p_ = 0.150), and the emotional valence of the image shown earlier (within in the tested pair) modulated temporal distance (B = 0.065 (SE = 0.019), F = 12.283, p < 0.001, η^2^_p_ = 0.110), which was again qualified by the interaction of emotional valence between the first and second images from the tested pair (B = −0.134 (SE = 0.024), F = 32.591, p < 0.001, η^2^_p_ = 0.26. Specifically, temporal distances continued to be remembered as significantly larger following [Neu-to-Neg] transitions compared to all other transitions ([Neu-to-Neu] (p < 0.001, *d_z_* = 0.775), [Neg-to-Neg] (p < 0.001, *d_z_* = 0.923) and [Neg-to-Neu] (p = 0.006, *d_z_* = 0.373). Finally, temporal distances for negative images within a negative event continued to be remembered as shorter than distances for neutral images processed within neutral events (p=0.017, *d_z_* = 0.328).

### Negative images encoded later in a sequence may boost temporal order memory

Next, we examined whether perceptually induced event boundaries produced the oft-observed cost in temporal order accuracy compared to within-boundary events (Clewett & Davachi, 2021; Heusser et al., 2018). We observed a trend-level effect of border color change on temporal order memory, such that temporal order memory was numerically lower for items sampled across (compared to within) border color changes (B = −0.057 (SE = 0.031), χ^2^_(1)_ = 2.94, *p* = 0.086, OR = 1.059, BF= 3.13) (**Figure 3A**). In addition, we found that the emotional valence of the more recently-encoded image from the tested pair modulated temporal order accuracy, such that order memory accuracy was higher for pairs that included a recently-viewed negative (compared to neutral) image (B = 0.110 (SE = 0.052), χ^2^_(1)_ = 4.49, *p* = 0.034, OR = 1.116, BF= 6.17). The three-way interaction of border color, first and second valence on temporal order accuracy was not significant (B = 0.049 (SE = 0.060), χ^2^_(1)_ = 0.68, p = 0.409, OR = 1.050, BF = 1.05). Other factors, including first-image emotional valence (B = 0.006 (SE = 0.054), χ^2^_(1)_ = 0.02, *p* = 0.902, OR = 1.006, BF= 0.85), or interactions amongst these factors, did not reach statistical significance, *p*s > 0.4 (For full model results, including Bayes Factors, means and SEM per condition, see **Supplementary Figure 3** and **Supplementary Tables 3 and 4**).

**Figure 3.**
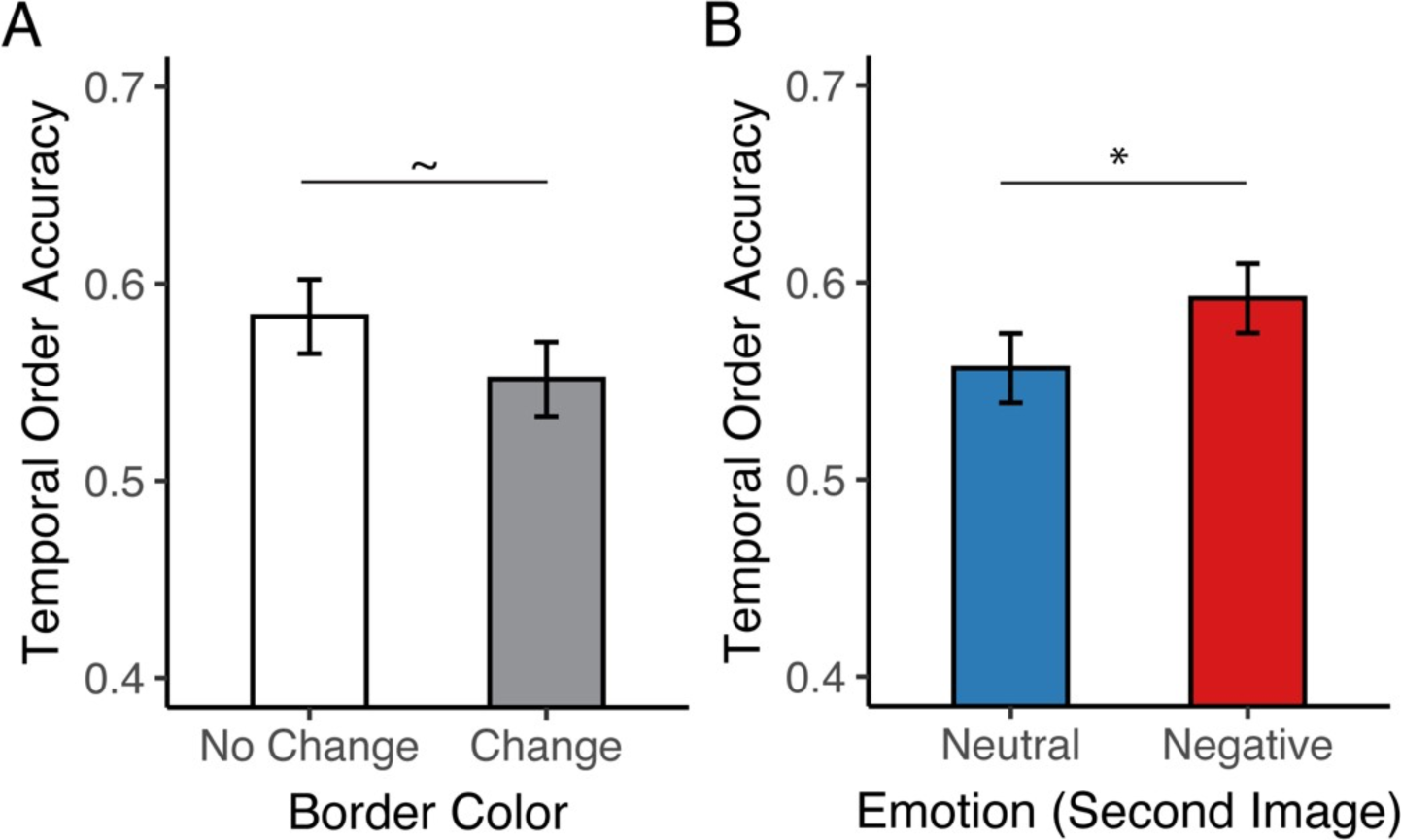
Impact of emotion and emotional-event transitions on temporal order memory. **(A)** Temporal order memory accuracy tended to be reduced for pairs of items sampled across (vs. within) border color changes. **(B)** Temporal order accuracy was significantly higher for image pairs that included a recently-viewed negative (vs. neutral) image. Error bars: within-subjects 95% confidence interval for each condition, computed using Morey’s (2008) method. \**p* < 0.05. ∼*p* < 0.1.

#### Control Analysis: item recency

Given that the sequence position of images in a tested pair itself may influence temporal order memory accuracy (Bjork & Whitten, 1974; Singh & Howard, 2017; Yntema & Trask, 1963), we included the sequence position of tested image pairs in our model to examine whether it exerted an effect. We found that image-pair sequence position was not a significant predictor of temporal order accuracy in our study (B = −0.021 (SE = 0.013), χ^2^ _(1)_ = 2.52, *p* = 0.113, OR = 0.979); moreover, the main effect of border color change (B = −0.061 (SE = 0.031), χ^2^ _(1)_ = 3.29, *p* = 0.070, OR = 0.941) and valence of the later-presented image (B = 0.110 (SE = 0.052), χ^2^ _(1)_ = 4.50, *p* = 0.034, OR = 1.117) remained consistent with results from the original model after accounting for sequence position.

### Emotion-driven shifts on temporal distance memory correlates with dispositional negativity

Next, we examined whether the magnitude of the impact of emotional-state shifts on temporal memory was associated with trait variation relevant to psychopathology vulnerability. To that end, we tested whether dispositional negativity, a latent factor obtained from trait mood and anxiety symptomatology, was associated with temporal memory dilation produced by shifts from Neutral-to-Negative events. To examine this dilation score while controlling for inter-individual variation attributable to other non-specific effects, we performed a residualization procedure controlling for variation in the reciprocal score (Negative-to-Neutral) across individuals (see *Methods*). We found that dispositional negativity was positively correlated with temporal memory dilation produced by the onset of negative events (r_(78)_ = 0.255, *p* = 0.023, **Figure 4**) ^4,5^. In contrast, dispositional negativity was not associated with variation in temporal memory following Negative-to-Neutral shifts (r_(78)_ = −0.080, *p* = 0.483). The difference between these correlation coefficients was at the trend-level, such as the correlation of dispositional negativity and this dilation effect was numerically higher than the correlation with remembered temporal distance following Negative-to-Neutral shifts (z = 1.895, *p* = 0.058).

**Figure 4.**
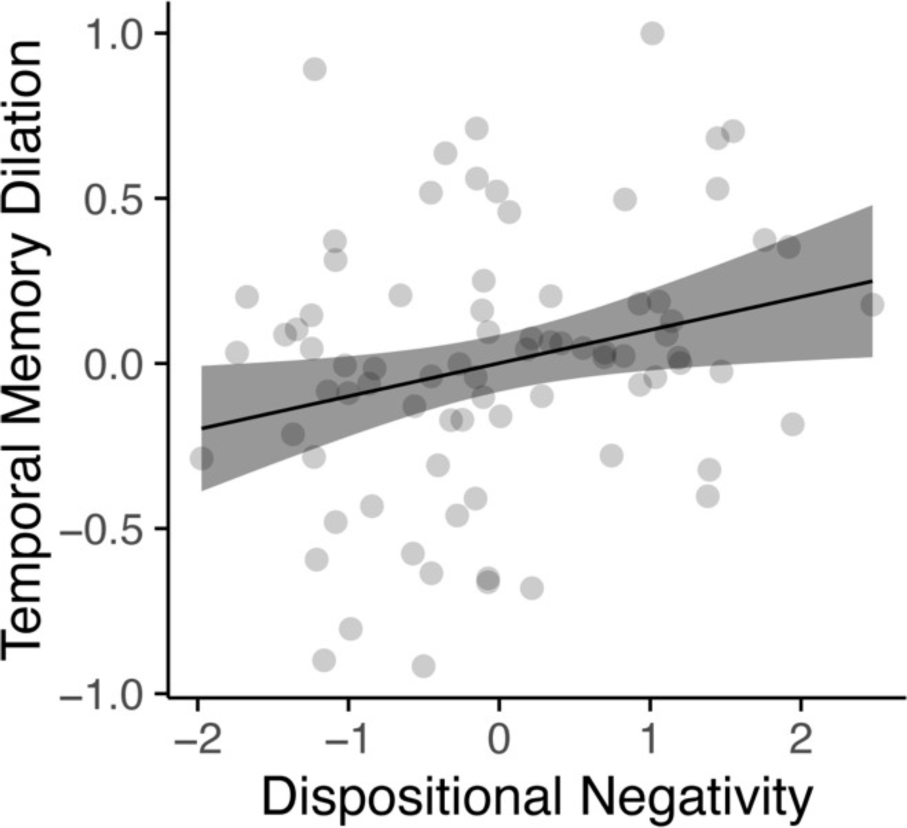
The association between temporal memory and dispositional negativity. Greater temporal memory dilation produced by neutral-to-negative emotional shifts was associated with higher dispositional negativity, *r*(78) = .255, *p* = .023.

In summary, increases in subjectively remembered temporal distances produced by the onset of negative emotional events were associated with trait variation in emotionality, or affective style (Davidson, 2000; Lapate & Heller, 2020).

### Dissociation of the emotional modulation of temporal distance versus order memory

Taken together, our results suggest distinct effects of emotional-state shifts on temporal distance and order memory. To quantify the potential dissociation between emotion-driven changes in temporal distance versus temporal order memory in our paradigm, we next examined whether emotion-driven shifts in these two temporal memory metrics were correlated across subjects. To do so, we examined the association between delta scores indexing emotion-dependent changes in temporal distance and temporal order accuracy for all condition pairs (see *Methods*). We found that temporal distance and order accuracy were largely *uncorrelated* across subjects (all *p*s > 0.09, **Table 1**), suggesting that these two temporal memory metrics dissociate, at least following dynamic emotional changes.

## Discussion

Fluctuations in internal emotional states are exceedingly common in everyday life. Yet, we are only beginning to understand precisely how emotional states and emotional-state dynamics modulate temporal memory, which is well known to be impacted by perceptual and semantic contextual shifts (Clewett & Davachi, 2021; McClay et al., 2023; Palombo & Cocquyt, 2020; Petrucci & Palombo, 2021; Radvansky & Zacks, 2017; Shin & DuBrow, 2021; Talmi et al., 2019; J. Wang et al., 2022; Zacks, 2020). Here, we designed a novel Emotion Boundary Task that orthogonally manipulated the valence and border color of emotional pictures shown in sequences. This experimental design yielded distinct emotion boundary conditions, which varied in the direction emotional-state shifts (Neutral-to-Negative and Negative-to-Neutral). We measured temporal memory—including subjectively remembered temporal distance and temporal order memory—for images sampled from within and across emotional events. First, we found that negative images encoded within negative event sequences were remembered as closer in time than neutral images encoded within neutral-event sequences. Second, temporal distance was remembered as longer for items spanning across Neutral-to-Negative valence transitions (compared to all other conditions)—indicating that Neutral-to-Negative emotional event shifts produced robust temporal dilation in memory. Moreover, the magnitude of this negative-emotion driven temporal memory dilation correlated with trait dispositional negativity, suggesting the relevance of emotion-temporal memory interactions for adaptive emotional functioning. Of note, these effects were consistent across changes of perceptual context (i.e., border color)—and specific to temporal distance (and not temporal order) memory. Collectively, these results suggest that emotional events processed within and across emotional state transitions produce distinct impacts on temporal memory.

### Sequential negative emotional events produced temporal compression in memory

We found that negative images encoded within negative-emotional sequences were judged as closer in time compared to images from neutral sequences, which we speculate may be due to an increased representational similarity of temporal contexts linking negative compared to neutral items, potentially resulting in increased event integration in memory (Ezzyat & Davachi, 2011; Zacks et al., 2007). Prior studies using free recall metrics showed that negative items appear to cluster together in retrieval, altering otherwise chronologically organized free recall (Barnacle et al., 2016; Long et al., 2015; Talmi et al., 2007). Recent computational models indicate that this emotional clustering phenomenon may result from a shared emotional context by negative items that strengthens their binding compared to neutral items (Cohen & Kahana, 2022; Talmi et al., 2019)—suggesting that negative emotional items may share greater representational similarity (compared to neutral ones) (Riberto et al., 2019) (see also: *‘Interconnectedness between semantic similarity, emotional valence, and temporal distance’* section below). Collectively, these recent findings are consistent with a potential distortion of the representation of temporal contexts for emotion-relevant items that surround emotionally evocative events.

Relatedly, we recently proposed that function of amygdala-hippocampal and entorhinal circuitry may underlie temporal context compression following the onset of negative emotional experiences (J. Wang et al., 2022). The granularity of temporal context encoding by hippocampal-entorhinal circuitry has been shown to track the temporal distance between events, with greater neural pattern similarity typically linked to shorter subjectively remembered (or actual) temporal distances (Deuker et al., 2016; DuBrow & Davachi, 2014; Ezzyat & Davachi, 2014; Folkerts et al., 2018; Lositsky et al., 2016; Nielson et al., 2015; Thavabalasingam et al., 2019). The amygdala, a set of nuclei involved in the early appraisal of emotionally meaningful stimuli, projects strongly to the hippocampus (A. K. Anderson & Phelps, 2001; LeDoux, 2007; Phelps & LeDoux, 2005; J. Wang & Barbas, 2018). Recently-unveiled molecular and anatomical features of those amygdala-hippocampal projections suggest that amygdalar inputs may compete with temporal signals in the hippocampal-entorhinal circuitry after the onset of high-arousal negative events, resulting in poorer-fidelity coding of subsequent time points (Tambini et al., 2017; J. Wang et al., 2022; J. Wang & Barbas, 2018; Yang et al., 2016; Zhou et al., 2009). Therefore, the relative temporal compression observed for sequentially presented negative items in our study may result in part from amygdala-elicited distortion of temporal coding in hippocampal-entorhinal circuitry, culminating in greater neural activity pattern similarity of temporal contexts shared by sequential negative items (versus sequential neutral items). Future neuroimaging work detailing the representational structure of temporal contexts sampled from within and across negative and neutral events will be required to fully examine this conjecture.

### Shifting from neutral to negative emotional states dilates time in memory

Despite fixed image presentation intervals, we found that the time interval between neutral and negative events was remembered as longer than the time elapsed between Negative-to-Neutral event transitions. This subjectively dilated time associated with Neutral-to-Negative valence shifts was also remembered as longer than the time elapsed between sequentially presented neutral or negative images. Consistently, a recent study using musical pieces to manipulate emotion during neutral-object encoding found that music-evoked emotional shifts toward a relatively more negative state produced larger remembered temporal distances for objects encoded during those emotional transitions (McClay et al., 2023).

Regarding the putative underlying mechanisms that may explain this emotionally ‘asymmetric’ boundary and resulting temporal dilation when shifting from neutral to negative events, we believe that a mixture of attentional and prediction-error factors are likely at play. First, event boundaries are often experienced following abrupt changes in conceptual (e.g., agent’s goal) and/or perceptual (e.g. visual and spatial) features, which engage additional attentional resources (Baker & Levin, 2015b, 2015a; Huff et al., 2012; Radvansky & Zacks, 2017), autonomic arousal (Clewett et al., 2020; Kurby & Zacks, 2008), and increase prediction errors (DuBrow et al., 2017; Reynolds et al., 2007; Shin & DuBrow, 2021; Zacks et al., 2007, 2011), which may lead to updating one’s model of the current environment (Reynolds et al., 2007; Zacks et al., 2007). This model updating is thought to underlie event segmentation in memory (Ezzyat & Davachi, 2011), which can increase the remembered temporal distance between boundary-spanning items, putatively due to reduced neural pattern similarity in the hippocampus across event boundaries (Ezzyat & Davachi, 2014). Similarly, a wealth of prior electrophysiological and behavioral data suggests that negative events engage greater attention (Bennion et al., 2013; Delplanque et al., 2006; Dewhurst & Parry, 2000; Kamp et al., 2012; Mather, 2007; Mather & Sutherland, 2011) and elicit greater changes in autonomic nervous systems (Bradley et al., 2008; Krebs et al., 2018; Snowden et al., 2016) compared to neutral events, in part due to amygdala engagement and resulting downstream and upstream modulation (Adolphs, 2008; Mather & Sutherland, 2011). Neutral-to-negative event transitions have been argued to produce greater prediction errors compared to sequential neutral or negative events (Astikainen & Hietanen, 2009; Chang et al., 2010; Kimura et al., 2011; Zhao & Li, 2006). Thus, acute neutral-to-negative transitions may change an otherwise cohesive mental model of the environment into a distinct model, resulting in temporal dilation immediately following the shift into a more negative state. In sum, it is possible that mechanisms previously implicated in event segmentation of relativelyneutral events may be preferentially engaged when experiencing neutral-to-negative state transitions, which may modulate temporal coding in the hippocampus, and increase subjectively experienced and remembered time.

### Interconnectedness between emotional valence, semantic similarity, and temporal distance

It is important to point out that differences in semantic similarity between negative and neutral visual events can partially account for temporal compression (and/or clustering effects) observed in response to negative (vs. neutral) emotional stimuli. Indeed, using an ANN-derived proxy for semantic similarity, we found that negative images were associated on average with higher semantic similarity estimates (compared to neutral images). This finding aligns with previous studies showing that negative words and pictures are often rated as having higher semantic relatedness compared to neutral ones (Gallo et al., 2009; Riberto et al., 2022; Talmi & Moscovitch, 2004; reviewed in Riberto et al., 2019). Critically, despite these effects underscoring the interconnectedness of semantic similarity, emotional valence, and temporal distance, we found that our temporal distance memory results held even after controlling for ANN-derived semantic similarity estimates. Specifically, temporal distances continued to be remembered as larger following [Neu-to-Neg] shifts compared to [Neg-to-Neu]—and smaller when image pairs were drawn from within negative (vs. neutral) events—even after controlling for differences in semantic similarity. This important control analysis suggests that emotional-event changes sculpt memory for temporal distance between those events beyond the (also robust) effects that semantic similarity can exert on temporal memory. Collectively, these results underscore the importance of controlling for semantic similarity in future studies of emotion and temporal memory—using estimates derived from behavioral judgments and/or existing ANNs tools—to fully disentangle the extent to which emotional effects in temporal memory are emotion-specific, or may stem from interrelated semantic factors.

### Temporal distance and temporal order judgments are differentially influenced by emotion

We found a distinct modulation of emotional states and state transitions on remembered temporal *distances* compared to temporal *order* memory, suggesting a dissociation between them. Results from a recent event boundary study suggest that temporal order effects may be more pliable and susceptible to paradigm variation compared to temporal distance memory. In their study, Wen and Egner found that event boundary saliency as well as the availability of a previous encoding context during retrieval substantially impacted whether and how temporal order memory was modulated by event boundaries (Wen & Egner, 2022). In contrast, temporal distance memory showed consistent event boundary effects (i.e., temporal distance dilation) across these manipulations (Wen & Egner, 2022). It is also possible that the temporal distance memory task is more sensitive to changes in emotional context compared to the temporal order task. For example, given the well-established modulation of attention by emotional items, participants may feel that they process emotional stimuli for longer (Johnson et al., 2019), thereby attenuating the “psychological temporal interval” that is experienced until the next item is shown, and producing shorter remembered temporal distances for Negative-to-Neutral, compared to Neutral-to-Negative conditions. In addition, the dissociation observed in our study between temporal order and distance memory may also arise partly from the fact that they represent distinct types of judgments. While temporal order is an objective judgment; the temporal distance metric reflects a subjective judgment. Neurally, while hippocampal coding appears to be involved in supporting both temporal distance and temporal order memory (Bellmund et al., 2020; DuBrow & Davachi, 2017), additional mechanisms have been postulated to support temporal order judgments that may or may not contribute to temporal distance estimates. Those include, for instance, theta gamma coupling (Heusser et al., 2016), and perirhinal-supported item-strength signals (DuBrow & Davachi, 2017; Jenkins & Ranganath, 2016), which, if confirmed to preferentially guide temporal order versus distance judgments, may jointly contribute for this apparent cacophony of results found when examining temporal order (versus temporal distance) memory.

Of note, temporal order memory accuracy was on average higher for image pairs containing a recently-shown negative (compared to neutral) image. On the one hand, high arousal negative emotion has been found to increase temporal order accuracy when processed in a naturalistic environment (Dev et al., 2022), which agrees with this result. Prior investigators have suggested that relative item strength (and/or saliency differences between items) can help support performance in temporal order tasks, such as the one adopted in our study (DuBrow & Davachi, 2017; Jenkins & Ranganath, 2016). Given that item memory for negative stimuli is often enhanced compared to memory for neutral stimuli (Bowen et al., 2018; Cahill & McGaugh, 1995; Kensinger & Corkin, 2003; LaBar & Cabeza, 2006; Ritchey et al., 2008), it is possible that our finding of higher temporal order memory accuracy for image pairs that contained a recently-shown *negative* (compared to neutral) image stems in part from the greater saliency of negative emotional items, which may provide a “boost” to temporal order memory accuracy (DuBrow & Davachi, 2017; Jenkins & Ranganath, 2016). On the other hand, a temporal-order memory enhancing impact of negative emotional images contrasts with a recent finding indicating that music-evoked transitions to a more negative emotional state reduces temporal order accuracy for neutral objects (McClay et al., 2023). It is possible that the emotional modulation of temporal memory varies by whether temporal memory is measured for items that were *themselves* emotionally evocative versus extraneous to the emotion-inducing stimuli (Mather, 2007; McClay et al., 2023); or, in other words, that negative emotion may differentially impact temporal memory for negative events compared to neutral items processed during negative states (McClay et al., 2023). Nonetheless, because our finding of higher temporal order memory accuracy for image pairs containing a recently-shown negative (vs. neutral) image was not a-priori predicted, it will be important to replicate it in the future. Moving forward, studies measuring item and temporal memory for events varying in emotional-episode relevance will be required fully dissect the factors driving the seemingly inconsistent impact of negative emotion on temporal order memory across studies.

### Inter-individual variation in temporal memory dilation after the onset of negative events correlates with dispositional negativity

Proneness to experience temporally persistent negative affect is a central feature of mood and anxiety disorders (Davidson, 2015; Heller et al., 2009; Kuppens & Verduyn, 2017; Lapate et al., 2014). In addition, individuals suffering from post-traumatic stress disorder (PTSD) and depression often report recurrent and intrusive aversive memories, suggesting potential alterations in the representation of temporal context surrounding aversive events (Davidson, 2000; Ehlers & Clark, 2000; LeMoult & Gotlib, 2019). Indeed, recent computational models suggest that interactions between emotional and temporal context coding may result in temporally persistent negative experiences in mood disorders (Cohen & Kahana, 2022). Consistent with these empirical findings and computational models, we found a positive association between the extent of the subjective dilation of time in memory following Neutral-to-Negative transitions and trait dispositional negativity, a known risk factor for mood and anxiety disorders (Shackman et al., 2016). In the future, it will be important to determine whether this finding may be due to increased attentional biases toward negative information (also known to be altered in mood and anxiety disorders (Bourke et al., 2010; Hamilton et al., 2012; Mennen et al., 2019; Mogg et al., 1995; Norris, 2021)—or, alternatively, whether it may stem specifically from less robust temporal context encoding or maintenance in at-risk individuals, which could precede the onset of negative events (Li & Lapate, 2023; J. Wang et al., 2022).

### Limitations and future research

The following limitations warrant additional investigation. First, the emotional manipulation and emotional-event transitions in the current study included only negative and neutral events. Understanding how positive emotions interact with temporal memory during dynamic emotional state transitions is a burgeoning area of research (Clewett & Davachi, 2021; Li & Lapate, 2023; McClay et al., 2023) that is essential for a full understanding of emotion-temporal memory interactions, as it permits isolating negative-valence specific from emotional arousal effects. Relatedly, in contrast to our a-priori hypothesis, we did not find that temporal distance judgments of Negative-to-Neutral transitions was compressed. However, the lack of positively valenced events (and/or Positive-to-Neutral transitions) precluded us from being able to characterize whether Negative-to-Neutral transitions may produce relative compression compared to a similarly arousing (and attentionally engaging) control condition. Second, our study aimed to induce strong emotional-state shifts to and from negative states, and therefore selected negative images low in valence, which are inherently higher on arousal than neutral images. Because arousal has been shown to modulate temporal order memory (Clewett & Davachi, 2021; Dev et al., 2022), future work employing psychophysiological measures to track a wide range of arousal levels elicited by both negative and positive emotional events will be required to determine the specific roles of valence *and* arousal in sculpting temporal distance and order memory. Third, our study only tested temporal memory for adjacently encoded items. Studies have shown that the interval between tested items can modulate temporal order accuracy and temporal distance judgments – for instance, a shorter interval between tested stimuli produces lower temporal order accuracy (DuBrow & Davachi, 2013, 2014; Gurguryan et al., 2021; Wen & Egner, 2022). Fourth, and relatedly, it is theoretically possible that certain participants could become aware of the lack of interval variability between image pairs when the interval between them is fixed, potentially compromising the quality of behavioral data obtained in the temporal distance task. In our study, following prior work, we adopted a long (16-image) encoding sequence, followed by a 45-second distractor task, which collectively minimizes the possibility that participants would be able to maintain a high-quality, conscious recollection of sequential intervals and their variability (or lack thereof). Accordingly, participants’ objective accuracy in the temporal order accuracy, while above chance (50%), was overall low (57%) (Supplementary Table 3). Nonetheless, it will be important to replicate and extend the current insights regarding how emotional-state dynamics alter temporal in future studies that adopt variable intervals between emotional items in emotional-event sequences.

## Conclusion

Our findings demonstrate that shifting from neutral to negative emotional events dilates time in memory—and that the magnitude of this emotion-driven distortion in temporal memory is associated with individual differences in trait dispositional negativity. Collectively, these findings underscore the interconnectedness of emotion and temporal coding processes, and pave the way for future work detailing their import for adaptive emotional functioning.

## Supporting information

https://osf.io/zr7hx/

## Acknowledgments

The authors thank Mengsi Li, Joanne Stasiak, and Reicher Bergstein for helpful discussions and assistance with manuscript preparation, and Zishi Ding and Mia Jeffery for assistance with data collection.

## Footnotes

1 In the current and similar studies, *temporal dilation* is referred as an increase of remembered temporal distance between conditions—relative to the average temporal distance observed in the study and/or relative to the temporal distance observed in other conditions—whereas *temporal compression* refers to a relative *decrease* in temporal distance.

2 The emotional manipulation in the current study was anchored on normative ratings of emotional valence, which were used to select negative vs. neutral pictures—we refer to this dimension as ‘emotional-valence’ following the conventional nomenclature in IAPS studies (Lang et al., 2008; Lang & Bradley, 2007). However, note that negative emotional pictures that are low in valence inherently differ from neutral pictures in both valence and arousal dimensions.

3 Note that because the temporal intervals between the sampled images were fixed, changes in temporal distance estimates reflect changes in *subjectively remembered* time.

4 Due to the relatively low loading of ATQ scores in the originally planned factor analysis, we also calculated the correlation between temporal memory dilation and dispositional negativity factor scores obtained after excluding the ATQ. This analysis replicated our results, with higher dispositional negativity correlating positively with the magnitude of temporal memory dilation across individuals (r = 0.248, *p* = 0.026).

5 Note that similar results were obtained following residualization procedures using the other available conditions— namely, after residualizing temporal distance memory for [Neu-to-Neg] event transitions for [Neu-to-Neu] (r(78)=0.197, *p*=0.080) or [Neg-to-Neg] r(78) = 0.236, *p* = 0.035) transitions.

