## Supplementary material for "Emotional state dynamics impacts temporal memory": https://osf.io/zr7hx/

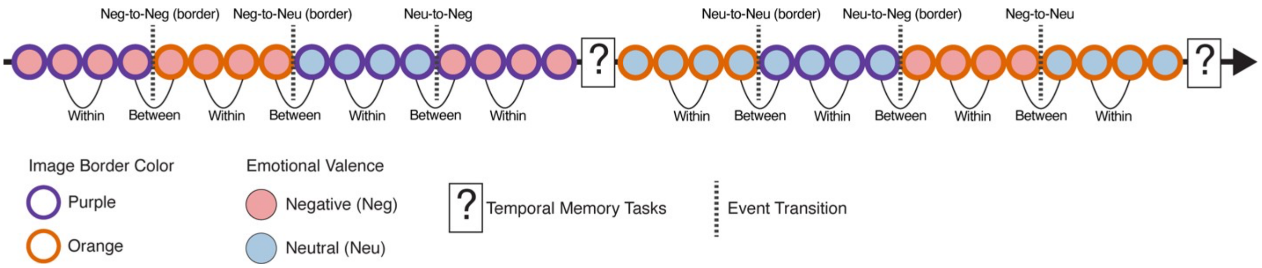


***Supplementary Figure 1:*** **Schematic of an example run depicting all event types and event-transition conditions.** In each run of the Emotion Boundary Task, participants studied two lists of image sequences, each followed by the distractor task and temporal memory tasks (depicted by the question mark). Each 'event' consisted of four images (negative: red, or neutral: blue) surrounded by a border (Purple or Orange). Each encoding list consistent of four events. The curved black lines linking two events at the bottom of the timeline indicate the image pairs tested in the temporal tasks. Whether a tested image pair belonged to the same vs. different event is indicated under the curved black lines as "within" and "between", respectively. All event transition types are depicted on the top row.

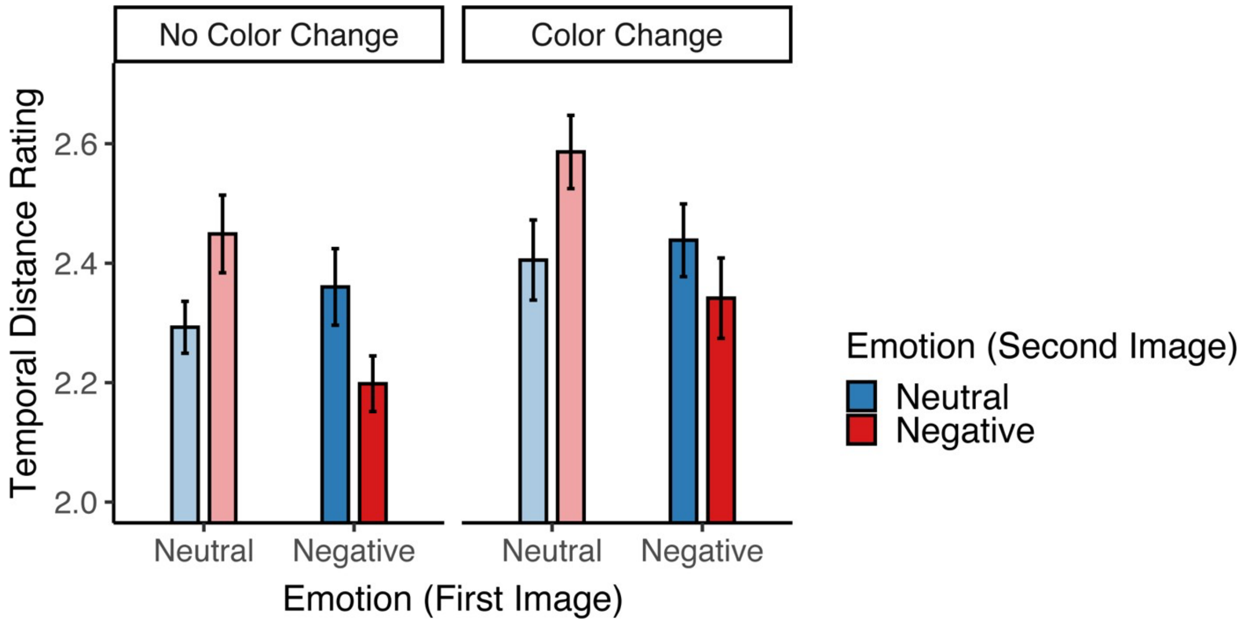


***Supplementary Figure 2.*** **Full model results for temporal distance memory.** Temporal distance ratings are plotted as a function of border color change and emotional valence of images encoded first and second within a tested image pair. Error bars: within-subjects 95% confidence interval for each condition.

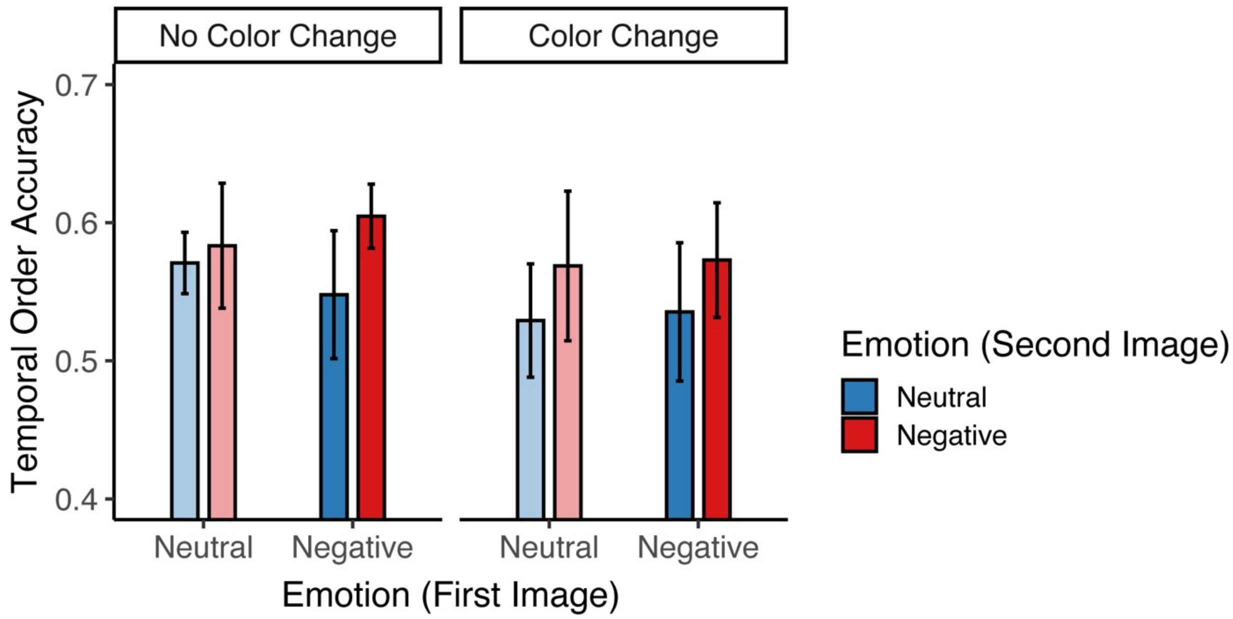


***Supplementary Figure 3.*** **Full model results for temporal order memory.** Temporal order accuracy is plotted as a function of border color change and emotional valence of images encoded first and second within a tested image pair. Error bars: within-subjects 95% confidence interval for each condition.

***Supplementary Table 1.*** **Mean and SEM for temporal distance ratings by condition.**

| **Condition** | **Mean** | **SEM** |
|---|---|---|
| Within Neutral | 2.293 | 0.022 |
| Within Negative | 2.198 | 0.023 |
| Border color change within neutral events | 2.405 | 0.034 |
| Border color change within negative events | 2.341 | 0.034 |
| Neutral-to-Negative (without border change) | 2.449 | 0.033 |
| Neutral-to-Negative (with border change) | 2.586 | 0.031 |
| Negative-to-Neutral (without border change) | 2.360 | 0.032 |
| Negative-to-Neutral (with border change) | 2.438 | 0.031 |

***Supplementary Table 2. Temporal Distance Results:* Full model results are shown, including Bayes factors for all main effects and interactions.**

| **Fixed effects** | **F** | ***p*** | **$\eta^2$** | **BF** |
|---|---|---|---|---|
| First image valence* | 14.630 | <0.001 | 0.140 | 678.910 |
| Second image valence | 0.669 | 0.415 | 0.007 | 1.25 |
| Border color change* | 27.119 | <0.001 | 0.170 | $2.933 \times 10^5$ |
| First_val × Second_val* | 40.568 | <0.001 | 0.31 | $1.943 \times 10^{11}$ |
| First_val × Change_color | 0.101 | 0.751 | $7.21 \times 10^{-4}$ | 0.13 |
| Second_val × Change_color | 1.021 | 0.314 | $8.11 \times 10^{-3}$ | 0.20 |
| First_val × Second_val × Change_color | 0.226 | 0.635 | $1.20 \times 10^{-3}$ | 0.06 |

First_val: First image valence; Second_val: Second image valence; Change_color: Border color change; BF: Bayes factor. * Denotes statistical significance at $p < 0.05$.

***Supplementary Table 3.*** **Mean and SEM of temporal order accuracy by condition.**

| Condition | Mean | SEM |
|---|---|---|
| Within Neutral* | 0.571 | 0.011 |
| Within Negative* | 0.604 | 0.012 |
| Border color change within neutral events~ | 0.529 | 0.021 |
| Border color change within negative events* | 0.573 | 0.022 |
| Neutral-to-Negative (without border change)* | 0.583 | 0.024 |
| Neutral-to-Negative (with border change)* | 0.569 | 0.028 |
| Negative-to-Neutral (without border change)* | 0.548 | 0.024 |
| Negative-to-Neutral (with border change)~ | 0.535 | 0.026 |

* Denotes statistical significance above chance (0.5) at $p < 0.05$. ~ Denotes numeric trend above chance (0.5) at $p < 0.1$.

***Supplementary Table 4. Temporal Order Results:*** **Full model results are shown, including Bayes factors for all main effects and interactions.**

| **Fixed effects** | $\chi^2$ | ***p*** | **OR** | **BF** |
|---|---|---|---|---|
| First image valence | 0.02 | 0.902 | 1.006 | 0.85 |
| Second image valence* | 4.49 | 0.034 | 1.116 | 6.17 |
| Border color change | 2.94 | 0.086 | 1.059 | 3.13 |
| First_val × Second_val | 0.58 | 0.446 | 1.039 | 1.10 |
| First_val × Change_color | 0.05 | 0.831 | 0.993 | 1.00 |
| Second_val × Change_color | 0.02 | 0.889 | 0.996 | 0.36 |
| First_val × Second_val × Change_color | 0.68 | 0.409 | 1.050 | 1.05 |

First_val: First image valence; Second_val: Second image valence; Change_color: Border color change; BF: Bayes factor. * Denotes statistical significance at $p < 0.05$.
